# Object detection through dynamic motor-sensory convergence

**DOI:** 10.1101/2025.07.16.665117

**Authors:** Guy Nelinger, Inbar Saraf-Sinik, Ehud Ahissar

## Abstract

To interact effectively with the world, animals coordinate how they move and how they sense. In natural settings, perceivers adapt their motor-sensory strategies as they approach and explore objects, dynamically shaping both the generation and interpretation of sensory cues. Previous attempts to explain this process by reducing it solely to neuronal representations have failed to capture the mechanisms underlying dynamic perception. In this study, we use precise behavioral tracking to investigate the initial phase of object interaction, asking: What motor-sensory strategies support object detection during natural approach? Using high resolution video tracking of rats freely exploring objects under infrared illumination, we analyzed how head and whisker dynamics evolved across sequential contacts. We found that whisker-object interactions converged toward a distinct line, α^*^, within a motor-sensory plane, where small changes in voluntary whisker movements produced large changes in whisker curvature, a sensory correlate of contact force. This convergence was actively controlled, predicted head movements and marked the completion of object approach. Convergence to α^*^ was consistent across different objects, suggesting it serves as an invariant motor-sensory contingency for object detection. Proximity to α^*^predicted the emergence of touch-induced pumps, a rapid motor-sensory reflex that further facilitated convergence within a single whisking cycle. Together, these results reveal that object detection is a closed-loop dynamic process, in which animals actively steer motor-sensory dynamics toward a robust detection-optimized state.

## Main

Natural environments are stochastic, rapidly changing, and present significant uncertainties. Yet, mammals are impressively effective at making sense of their surroundings and guiding behavior accordingly. They achieve this through generative motor-sensory processes, selecting the perceptual strategies that best serve their behavioral goals while adapting to environmental conditions^1–3^. For example, to judge a peach’s ripeness, broad and gentle pressure is typically applied; to find bruises, fingers sweep the surface as pressure is continuously adjusted. This kind of goal-dependent adaptation of motor-sensory strategy has been repeatedly demonstrated in whisker-based object localization. Object location can be encoded either through motor variables, sensory variables, or combined motor-sensory schemes, as demonstrated in head-fixed rats and biomechanical models^4–8^. This diversity is reflected in behavior, with studies in freely moving rats demonstrating that they actively employ and adapt different strategies depending on task demands^9–11^. Perception can therefore be supported by various coding schemes, and a range of strategies aimed at stabilizing motor actions, sensory signals, or their relations^2,3,12–15^. This behavioral flexibility means that sensory signals must be interpreted in light of the movements that generated them^16,17^ (we therefore refer to these combinations as motor-sensory, rather than sensorimotor).

Freely moving rats choose when and how to engage (or disengage) with the object of focus, exercising control over information flow and the level of perceptual accuracy. Over the course of exploration, control requirements change with instantaneous goals^9,18,19^; for example, object detection and localization, followed by focused exploration. Here, we examined such object-approach behavior in freely moving rats presented with a single object, and characterized the interaction between movements, object contact, and sensory signals. Studying rats that were free to select and control explorative motor variables, we asked: what motor-sensory contingencies support object detection? We found that over sequential contacts, the relation between changes in the whisker-base’s angle and curvature evolved towards a specific ratio which we term α^*^. We propose that proximity to α^*^represents, for the freely moving rat, the perceptual confidence of contacting an external object.

### Whisker-object contacts densely cluster around a distinct ratio in the motor-sensory space

Accurate quantification of motor and sensory measurements of the whisker was performed using high-spatio-temporal resolution videography (1024 × 1280 pixels, 500 frames per second, respectively; see **Methods**), under IR-only illumination. When whisking, rats actively control their whisker base-angle (θ; **Fig. 1a,b**). During whisker-object contacts, the whisker bends, producing changes in base-curvature (κ, which is inversely related to the radius of an osculating circle at the point, and thus measured in units of *m*^−1^, see **Methods**; **Fig. 1c**). Base-curvature during contact is proportional to the bending moment at the whisker base, which activates mechanoreceptors and consequently drives the trigeminal neurons^6,20,21^. We tracked the dynamics of motor variable, θ, and sensory variable, κ, for individual C-row whiskers bilaterally and examined how they varied with object contact. All other whisker-rows were trimmed to avoid occlusions, which hinder tracking (see **Methods**). Values of κ during non-contact whisking are not proportional to bending moment^22^, but are given as a reference.

**Fig. 1.**
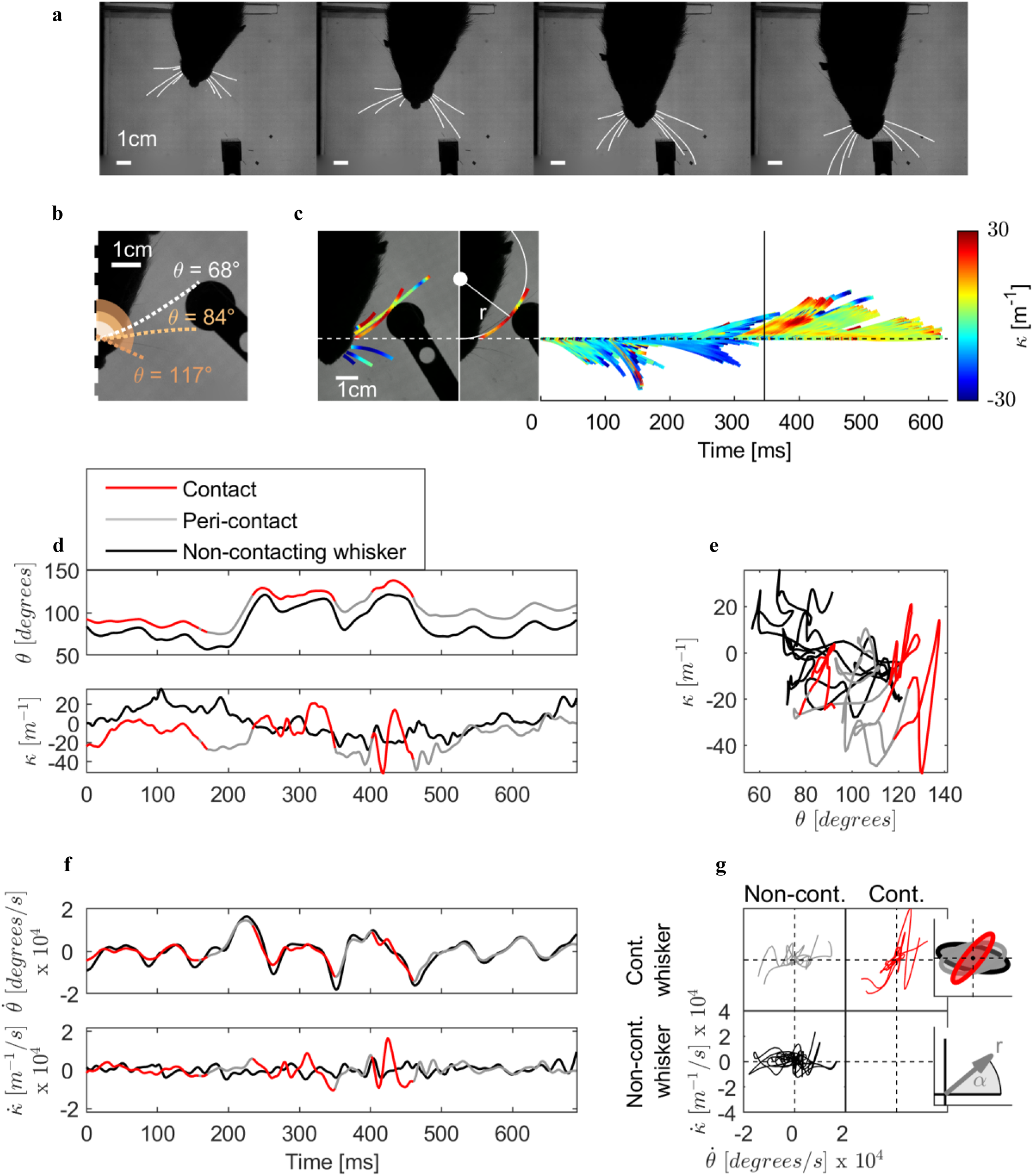
Motor and sensory variables of the whisker system during contact and non-contact. **a.** Four frames from a single trial. Data were acquired via top view videography and backlight IR illumination of freely moving rats. Whiskers were traced in each frame and tracked with consistent labeling across frames. **b.** The angle of the whisker’s base with respect to head-center axis is a motor variable termed base-angle and labeled θ. **c.** During contact, the curvature at the whisker’s base, labeled κ, is proportional to the bending moment, to which mechanoreceptors are sensitive. Left: curvature color-coded on the whisker traces. Middle: Same frame with one whisker singled out. Curvature at a point is proportional to the inverse radius of an osculating circle, thus expressed in units of *m*^−1^. Right: curvature of the same whisker over time, color-coded on its trace. Horizontal line marks the time of the frame shown to the left. Vertical dashed line denotes a follicle-aligned position. **d.** Example temporal dynamics of θ and κ, for two whiskers on the same side of the snout, tracked simultaneously. One whisker contacts an object (red, and gray for instances of non-contact), the other does not (black). **e.** Same data as in **d**, plotted on the (*θ*, κ) motor-sensory plane. **f.** Same as **d**, but for the time-derivatives of the variables, *θ̇* and *κ̇* Shown lowpass-filtered for visualization; unfiltered elsewhere unless otherwise noted. **g.** Contact and non-contact trajectories on the (*θ̇*, *κ̇*) phase-plane. Non-contact trajectories tend to evolve horizontally; contact trajectories tend to cluster around a distinct angle with respect to the origin. Top inset: shaded overlay of the trajectories. Ellipse orientation was obtained by transforming data to polar coordinates and minimizing the median absolute deviation (MAD) of their angular component; axis lengths reflect the MAD of angles around them. Bottom inset: The polar-plane equivalent to Cartesian (*θ̇*, *κ̇*) is termed (α, *r*).

As rats approached the object, they engaged in free-air whisking until making contact, usually unilaterally. Throughout the approach, rats typically varied whisker angle, inducing corresponding torsion-related changes in curvature^6,23^. In a representative example, we examined dynamics of two whiskers on the same side of the snout—one that made contact, and one that did not. Angle dynamics were similar across whiskers (**Fig. 1d**, top), but contact was associated with sharp transients in curvature. Nonetheless, the range of curvature values measured during contact still overlapped substantially with values measured in non-contact periods, both within the contacting whisker (**Fig. 1d**, bottom, gray) and compared to the non-contacting whisker (**Fig. 1d**, bottom, black). Thus, neither angle nor curvature are likely to make a reliable indicator of contact. Previous work in anesthetized rats and models has shown that an object’s position was ambiguously encoded by either angle or curvature separately but reliably read out when both were considered jointly^4,5,7,8^. In line with this, plotting angle and curvature jointly on a motor-sensory phase-plane produced partial separation of the non-contacting and contacting whiskers (**Fig. 1e**). However, epochs of contact did not occupy a unique contiguous region but rather overlapped with non-contact epochs. The (θ, κ) plane thus does not support a reliable representation of contact epochs in freely moving rats.

Since perceptual systems are typically tuned to changes rather than absolute values, a principle captured by Weber’s law^7^, we examined how object contact is reflected by the change in angle and curvature, using their time-derivatives denoted *θ̇* and *κ̇* (see **Methods)**. Considered individually, neither angle change nor curvature change showed robust separation of contact from non-contact (**Fig. 1f,g)**. The angle-change trace for the contacting whisker was nearly identical to that of the non-contacting whisker (**Fig. 1f**, top). Curvature change was preferentially associated with sharp increases during contact, although similar increases sometimes appeared in the non-contacting whisker as well (**Fig. 1f**, bottom). When examined jointly on a motor-sensory phase-plane, however, the joint dynamics of angle-curvature changes revealed a distinct region associated with contacts (**Fig. 1g**). Whereas non-contact trajectories typically evolved horizontally and covered much of the plane, contact trajectories closely traced a tilted line through the origin with a steep positive slope (**Fig. 1g**). In other words, instances of contact were associated with a specific ratio such that small changes in whisker angle induce large changes in curvature.

Therefore, we asked which combinations of the motor and sensory variables are statistically associated with object contact across the entire dataset (*n*_*whiskers*× *frames*_ = 616,708). To automatically classify data points as contact versus non-contact, we calculated the minimal distance between each whisker trace and the object’s surface. For this analysis, we excluded snout-object contact frames (see **Methods)**. The resulting distribution featured a pronounced left tail with a distinct mode at very small distances. This mode was used to set a data-driven threshold separating contacts from non-contacts at ∼1 mm (**Fig. 2a**; see **Methods**). Next, we plotted visit-rate maps of motor-sensory combinations on a polar plane, equivalent to the original Cartesian plane of angle-curvature changes (*θ̇*, *κ̇*). In this polar representation, angles, which we term α, correspond to specific ratios of curvature change to angle change; different combinations composing the same ratio α are mapped along individual radii from the origin (termed *r*; **Fig. 1g**). The transformation to polar coordinates highlights the importance of motor-sensory ratios, while maintaining a one-to-one mapping with original Cartesian coordinates. Since units of angle and curvature have no a priori correspondence, curvature was re-scaled before transformation to polar coordinates in all following analyses (see **Methods**).

**Fig. 2.**
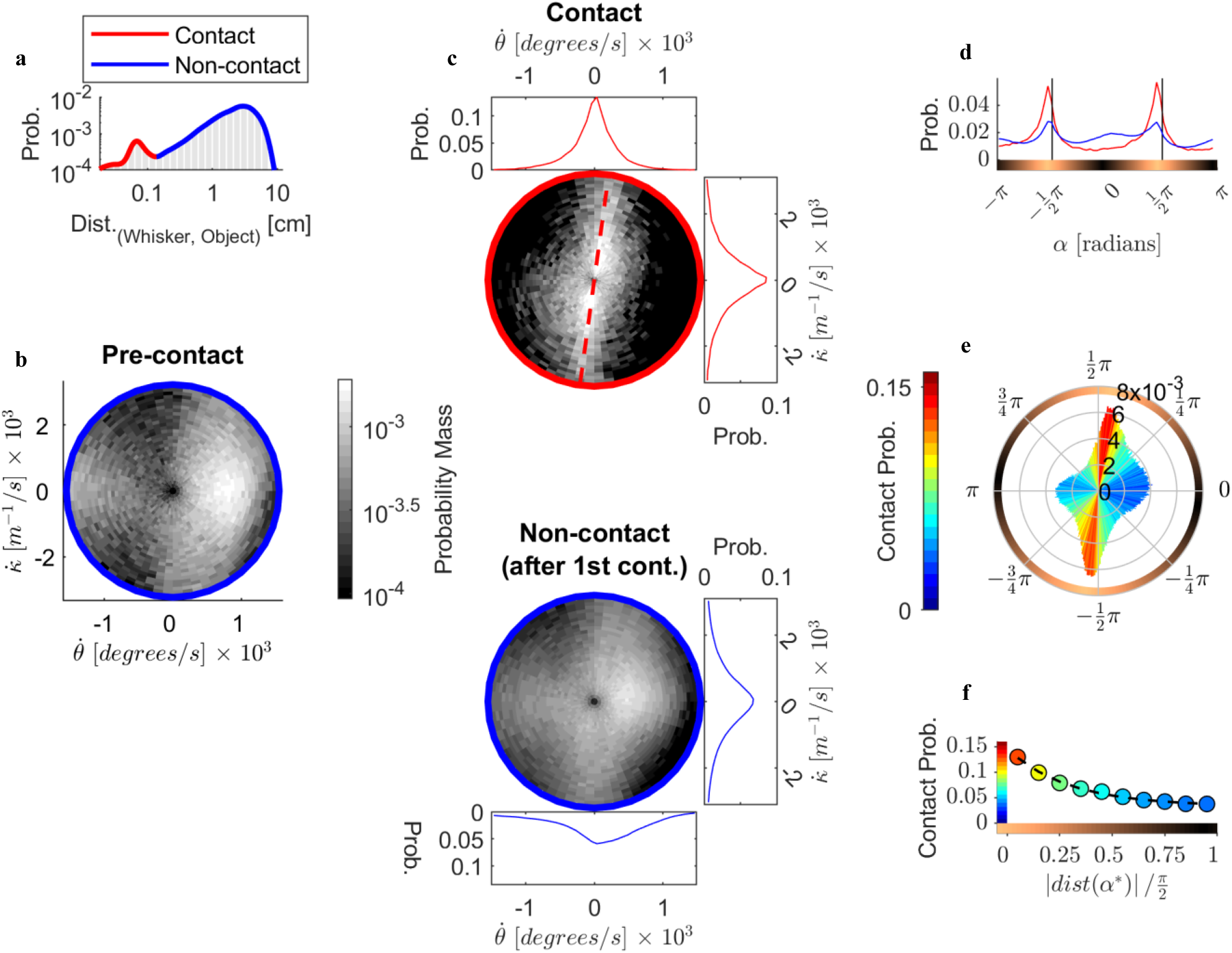
Whisker-object contacts are associated with a typical angle on the(*θ̇*, *κ̇*) plane. **a.** Whisker-object distance distribution across all data-points, excluding frames of snout-object contact (*n*_*whiskers*×*frames*_ = 616,708). Red and blue outlines denote contact and non-contact observations, respectively. **b.** Visit-rates on the (*θ̇*, *κ̇*)polar-plane prior to initial whisker object-contact (*n* = 120,486). **c.** Visit-rates on the (*θ̇*, *κ̇*) polar-plane following initial contact, separately for contacting (top, red; *n* = 44,621) and non-contacting whiskers after 1^st^ contact (bottom, blue, *n* = 451,601). Marginals show *θ̇* and *κ̇* Red dashed line marks the mode of (*θ̇*, *κ̇*) angles’ distribution in the polar phase-plane, labeled α^*^. The difference between contact and non-contact distributions was greater than the difference between pre-contact and non-contact distributions (permutation test on difference in Jensen-Shannon divergences, one-tailed, *p* < 0.001). **d.** Distribution of α, the angle with the origin of the (*θ̇*, *κ̇*) plane, for contacting (red) and all non-contacting (blue; same as in **a**) whiskers. The contact distribution had lower entropy (permutation test on entropy difference, one-tailed, *p* < 0.001). Light to dark brown color-bar denotes distance from α^*^. **e.** Bins of α colored by contact probability. Bar heights correspond to probability of α. Boundary circle color-code denotes distance from α^*^, as in **d**. **f.** Probability of contact as a function of the distance of α from α^*^. Black dashed trace denotes an exponential fit 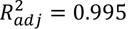 axis color-code denotes distance from α^*^, as in **d**., *p* < 10^−8^. X-axis color-code denotes distance from α*, as in **d**.

Before the first whisker-object contact occurred, combinations of angle-curvature changes covered much of the (*θ̇*, *κ̇*) plane, without a dominant peak (**Fig. 2b**, *n* = 120,486). Following the first contact, a pronounced difference emerged between contacting and non-contacting whiskers: contacts tightly clustered around a specific ratio, forming a line in the phase plane, which we label α^*^(**Fig. 2c**, top; *n* = 44,621; α^*^ denoted by the red dashed line, see **Methods**). In contrast, the non-contact distribution (after 1^st^ contact, **Fig. 2c**, bottom; *n* = 451,601) resembled the pre-contact distribution. The steep slope associated with α^*^ captures instances in which small angular changes induce relatively large changes in bending – as would be expected from a motor-sensory contingency supporting contact detection. The clustering around α^*^after initial contact was driven specifically by contact, rather than by a general behavioral change, as the difference between contact and non-contact (after 1^st^ contact) was greater than the difference between pre-contact and non-contact (permutation test using Jensen-Shannon divergence, *p* < 0.001; see **Methods**). Alternative phase-planes, expressing different combinations of angle, curvature, and their derivatives, produced less clear separation between contact and non-contact (**Extended Fig. 1**).

Focusing specifically on α, the ratio of angle-curvature changes, we found that its distribution was narrower during contact than during non-contact, with a sharp peak near α^*^(**Fig. 2d**; permutation test on Shannon entropy, *p* < 0.001, see **Methods**). Consistent with this, the probability of a single whisker making contact increased up to threefold near α^*^, relative to more distant values (**Fig. 2e,f**; exponential fit, 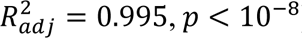). Taken together, these analyses suggest a strong association between contact occurrence and a distinct ratio of whisker angle-curvature changes, namely α^*^, which forms a plausible motor-sensory contingency representing contact.

### Motor-sensory dynamics converge towards the invariant contact ratio α^*^

Given a single sample, the probability of contact remains low even near α^*^. This fact, together with the use of multiple contact epochs, strongly suggests that our rats relied on accumulating evidence across contacts. To examine the dynamics of this evidence accumulation, we asked how α evolved during object approach (see **Methods**), and whether this differed between contact and non-contact whisking. We approximated the system’s flow separately for contact and non-contact observations, examining how changes in α depend on its current value. Flow was quantified using two attributes: mobility and directional tendency (see **Methods**). Mobility measures how large a change tends to be (calculated as *median*(|Δα|)). Directional tendency measures whether changes tend to evolve clockwise or counterclockwise (calculated as *p*(Δα > 0) − 0.5).

During retraction, dynamics were similar for contact and non-contact. In contrast, during protraction, contact and non-contact dynamics diverged: contact dynamics evolved away from α ≈ 0 and towards α^*^, whereas non-contact dynamics evolved away from α^*^and towards α ≈ 0 (**Fig. 3a,b**). This dissociation in directional tendency profiles highlights contact-induced effects on the flow of α: α^*^ is attractive only for contacts, and only during protraction—which is specifically relevant to exploration of the object ahead. Given the inherent asymmetry between protraction and retraction, α^*^ is resistant to perturbation only from one direction, i.e., it approximates a semi-stable attractor (see **Extended Fig. 2**).

**Fig. 3.**
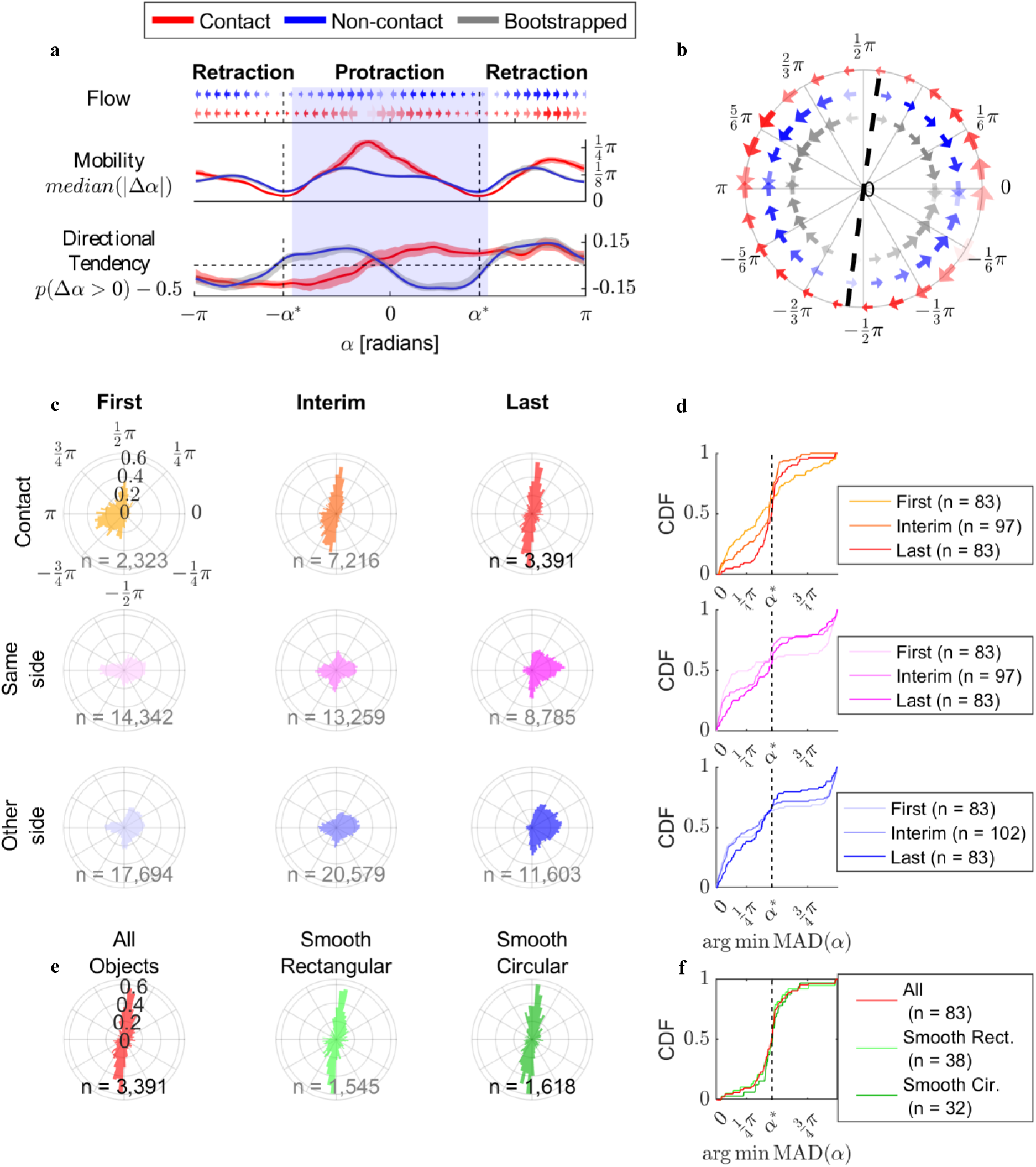
Contacts display convergence dynamics towards. α^*^**. a.** Flow dynamics of α during approach (Δα computed over 4ms intervals), for contact (red; *n* = 9,663) and non-contact observations (blue; *n* = 167,578). Top: arrow sizes denote mobility, shading indicates the strength of directional tendency. Middle: solid traces denote mobility, the absolute median step-size given current α. Shaded error bars denote a bootstrapped 95% confidence interval, with red sampled from the contact distribution, and gray from the combined distribution. Bottom: same as middle, but for directional tendency. **b.** Same data as in **a**, arrow sizes denote mobility, arrow shading denotes directional tendency. Gray arrows plot the mean of bootstrapped populations from panel **a**. **c.** Probability distributions of α values for *whiskers* × *frames* observations, grouped by cycles (in columns: first, interim and last; excluding trials with a single contact cycle), and by subset (rows: contact, same-side no contact, other-side no contact). Data are included from the start of the first contact cycle throughout approach. **d.** Same data as in panel **c**, but each cycle is summarized by the value that minimizes the median absolute deviation (MAD) for its subset of observations (computed modulo π). Traces show CDFs. The median of the last contact cycle distribution was indistinguishable from α^*^ (Wilcoxon signed-rank test, *W* = 1,547, *p* = 0.374). The variance of the last cycle distribution (darkest shades) for contact was smaller than that of non-contact on either the same side (Brown-Forsythe test, *F*_(1,164)_ = 31.752, *p* < 10^−7^) or the other side (*F*_(1,164)_ = 25.931, *p* < 10^−6^). **e**. Data from all last contact cycles (reproduced from top-right panel of **c**), shown combined for all three objects, and separately for two smooth objects differing only in shape. **f.** CDFs of the MAD-minimizing angle α for the data from panel **e**. α^*^did not significantly differ across all three objects presented (omnibus permutation test, *p* = 0.071). Specifically, no significant effect of shape was found between the smooth circular and smooth rectangular poles, which were presented in most trials (*p* = 0.78).

Are these fast, within-cycle, convergence dynamics towards α^*^ supported by slow dynamics across whisking cycles? For this analysis, we parsed the whisking trajectories into distinct cycles (see **Methods**) and included only trials in which a clear unilateral contact preference was demonstrated during the approach phase; otherwise, ordering of contact cycles became infeasible. To compare across trials with a different number of cycles, we grouped them into first, interim, or last - which ensured each trial contributed exactly one cycle to the first and last groups (*n*_*trials*_ = 83/155). Trials with a single-contact cycle are considered separately in the next subsection.

Collapsed across trials, contacting whiskers showed gradual convergence towards α^*^, from first to last cycles (**Fig. 3c**, top row). Conversely, non-contacting whiskers – both from the contacting side and the other side – remained broadly distributed across consecutive cycles (**Fig. 3c**, middle and bottom rows, respectively). For the non-contact groups, the last cycle distributions appear to cluster in the right hemi-plane because data only extend through the end of the approach phase, which typically ended before retraction began.

To quantify the observed pattern, we analyzed data at the cycle level by assigning each cycle with a single representative value, the angle α for which the median absolute deviation was minimized (MAD; see **Methods**). This ensured that longer cycles, or cycles where a larger proportion of the whiskers contacted the object, were not overrepresented, and that the first and last cycle groups were balanced. The resulting cycle-level statistics confirmed the convergence pattern described qualitatively above: for the distribution of last contact cycles, the median α aligned with α^*^ (**Fig. 3d**; Wilcoxon signed-rank test, *p* = 0.374), and its variance was narrower compared to that of simultaneous non-contact cycles, both ipsi- and contra-lateral to the contacting whiskers (Brown-Forsythe test, *p* < 10^−6^ for both comparisons). This convergence process is consistent with the dynamics of a controlled variable: a perceptually meaningful quantity that perceivers actively maintain near a desired value by adjusting their behavior^24^. Consistently, α^*^remained a reliable indicator of contact across all three objects (*p* = 0.071; omnibus permutation test), exhibiting almost identical behavior across the two objects that were presented in most of the trials and differed only in shape (**Fig. 3e,f**; *p* = 0.78).

### Correlation structures suggest **α** dynamics guide behavioral adaptation

The evidence above indicates that α converges towards α^*^over consecutive contacts during object approach. Such convergence is consistent with active control, but could also arise passively; for example, if rats follow a pre-planned sequence of body and whisker movements, where forward motion is pre-determined, and it is the diminishing distance to the object that gradually shapes the outcome of whisker-object contact. To distinguish these possibilities, we focused on trials in which approach included only one contact cycle (termed single-contact cycles, *n* = 26, see **Methods**), and compared these to the first and last contact cycles of multi-contact approaches. If α guides approach termination, it should converge on α^*^ even in single-contact trials, and these cycles should therefore more closely resemble last than first cycles. If convergence reflects passive constraints affecting both head and whiskers, a similar pattern should also be observed in other movement-related variables.

To evaluate whether α demonstrated the predicted pattern, we quantified its dynamics within each cycle using median absolute deviation from α^*^(*MAD*(α; α^*^)). As predicted, α’s values in single-contact cycles closely matched those observed in last cycles, but not first cycles, of multi-contact trials (**Fig. 4a**; permutation test on medians, first vs. single: *p* < 0.05; last vs. single: *p* > 0.05; FDR corrected across panels **a-d**). In contrast, approach velocity – defined per cycle as the mean change in head-object distance along the y-axis – differed in single-contact cycles from both first and last cycles (**Fig. 4b**; *p* < 0.05 for both comparisons). This challenges an interpretation in which α’s dynamics are merely a byproduct of pre-planned head-movement. We next asked whether this pattern of results was unique to α – or whether it may also be observed in other whisker-related variables. Specifically, we examined two additional variables: the radius on the polar-plane, termed *r*, representing the magnitude of motor-sensory change (quantified per-cycle by its 90^th^ percentile to capture trajectories’ peak, see **Fig. 1g**), and the shortest distance from whisker base to contact point, termed radial distance (RD; quantified per-cycle by its median). RD showed a similar pattern to α, with values in single-contact trials resembling last but not first contact cycles (**Fig. 4d**; first vs. single: *p* < 0.05; last vs. single: *p* > 0.05). In contrast, *r* did not exhibit the alignment observed in α and RD (**Fig. 4c**; *p* > 0.05 for both comparisons).

**Fig. 4.**
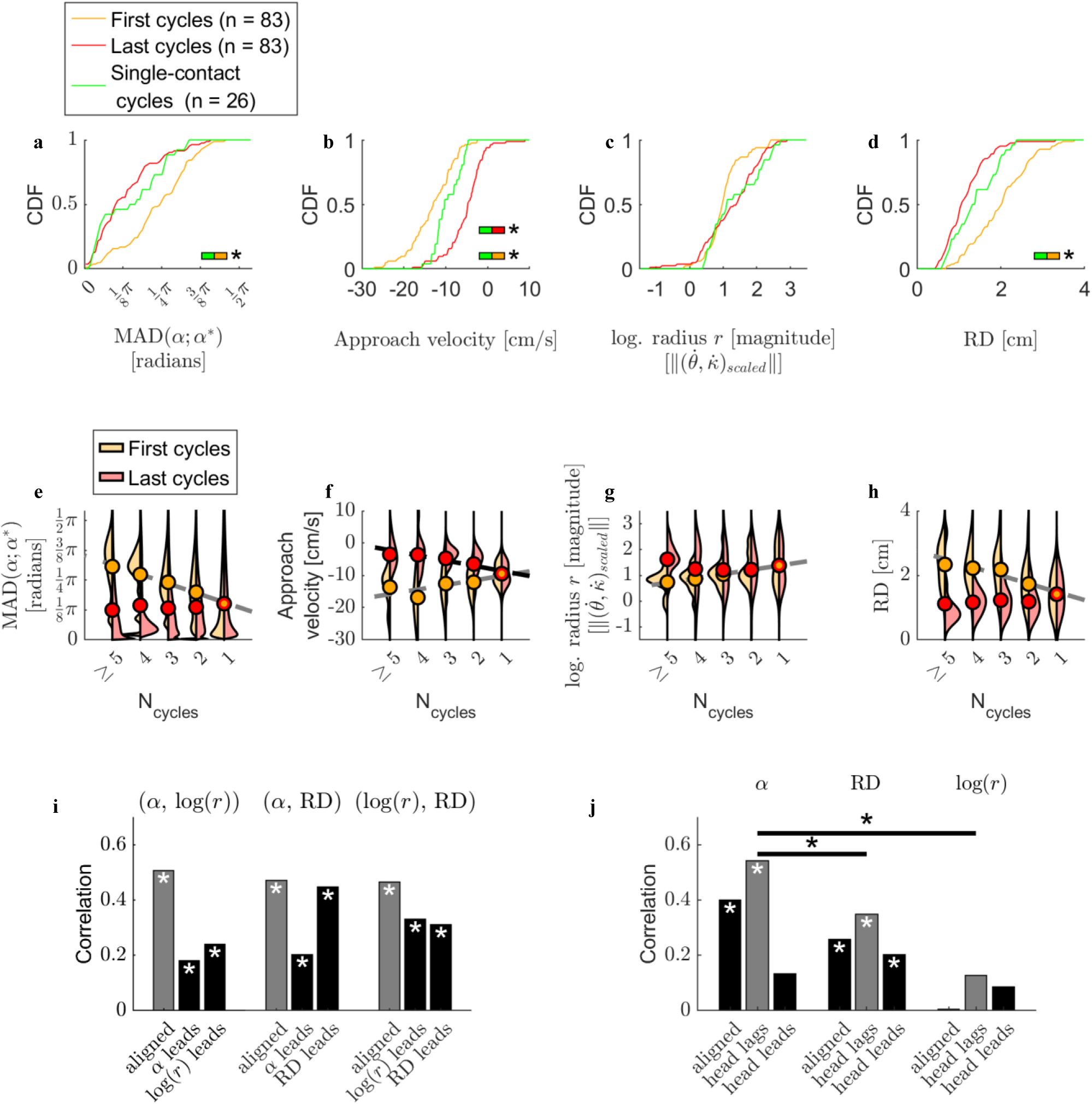
Correlation structures suggest α dynamics guide behavioral adaptation. **a–d**. CDFs of single-contact trials compared to those from first and last contact cycles in multi-contact trials. Split colored bars indicate significant differences between single-contact and first or last cycles (permutation test on medians, *p* < 0.05, FDR corrected across **a–d**). Variables, summarized per cycle: **a.** α, quantified as the median absolute deviation from α^*^; **b**. RD, the median radial distance from whisker follicle to contact point; **c.** *r*, the polar-plane radius, i.e. the magnitude 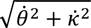 after scaling (see **Methods**), quantified as the 90^th^ percentile; **d**. Approach velocity, defined as the mean change in head–object distance along the y-axis. **e-h**. First and last contact values for each variable (same as in panels **a–d**), plotted separately as a function of the total number of contact cycles per trial. Violin plots denote kernel-density estimates; filled circles show means; dashed lines indicate significant linear fits (*p* < 0.05, FDR corrected across panels **e-h**); gray and black lines correspond to first and last contact values, respectively. **i.** Cycle-by-cycle Pearson correlations among the three whisker-related variables, α, log(*r*) and RD. For each pair of variables, observations are either aligned (gray bars, *n* = 289), or cycles are shifted such that one variable leads and the other lags (*n* = 180). White asterisks denote significance after FDR correction for all bars in the panel. **j.** Same as **i**, but for each whisker-related variable and approach velocity. Statistical comparisons were performed only on the aligned correlations, which showed the peak correlation values for each variable pair. Only the two pairs shown were tested. Horizontal bars denote significant differences in correlations (Fisher r-to-z transformation, *p* < 0.05, FDR corrected).

We expanded on this rationale to further explore which variables are under active control. If approach terminates when a controlled variable reaches some desired range, then trials with more contact cycles are expected to typically begin farther away from that range. In other words, the distribution of first contacts should vary depending on the number of following cycles, while the distribution of last contacts should be similar regardless of the number of preceding cycles. This analysis mostly agreed with the previous one, as α, RD, and here also *r*, all showed the expected pattern for controlled variables: values at first contact showed a linear dependency on the number of contact cycles, whereas values at last contact did not (**Fig. 4e,g,h**; *p* < 0.05 for significant slopes, FDR corrected across panels **e-h**). However, for *r*, variance increased from the first to the last cycle, as opposed to expectations from a controlled variable^24^. In contrast to these variables, approach velocity showed a linear dependency on the number of contact cycles in both first and last contacts, inconsistent with convergence toward a fixed target (**Fig. 4f**; *p* < 0.05). Finally, on a cycle-by-cycle basis, all three whisker-related variables were significantly correlated with one another (Pearson’s correlation coefficient, *r*_α,*r*_ = 0.508, *r*_α,*RD*_ = 0.471, *r*_*r*,*RD*_ = 0.466, *p* < 0.05 in all cases; **Fig. 4i**, gray bars). These correlations alone do not distinguish whether the variables co-vary or whether one drives the others, which motivated us to explore their temporal relationship.

To uncover a possible underlying causal relationship between these variables, we repeated the cycle-by-cycle correlation procedure. This time, however, we examined not only time-aligned correlations but also lagged the cycle-to-cycle correspondence by ±1 cycle to see whether any variable predicted another variable’s value in the following whisking cycle. All three whisker-related variables - α, *r* and RD - achieved their peak correlation with one another when time-aligned (i.e., at zero lag), suggesting that none of them drives the others (**Fig. 4i**, gray bars, *p* < 0.05, FDR corrected). In contrast, all three whisker-related variables predicted approach velocity in the next cycle better than in the current or previous cycle (i.e., peak correlation was when approach velocity lagged; **Fig. 4j**, gray bars). Approach velocity in the following cycle was best predicted by α (Pearson’s *r* = 0.542, *p* < 0.05, FDR corrected; see also **Extended Fig. 3**), less so by RD (Pearson’s *r* = 0.348, *p* < 0.05) and least by *r* (Pearson’s *r* = 0.129, *p* > 0.05). Among these head-lagged correlations, the relationship between approach velocity and α was significantly stronger than between approach velocity and either RD or *r* (Fisher r-to-z transformation, *p* < 0.05, FDR-corrected; black horizontal bars in **Fig. 4j**). Overall, these findings suggest that α’s convergence arises from selective adaptation, that it guides approach termination, and that its dynamics are broadly aligned with those of RD.

### Motor-sensory phase-plane dynamics guide the emergence of touch-induced pumps, which in turn further facilitate convergence

In some cycles, rats selectively exhibit a distinct whisking pattern: a partial retraction immediately followed by additional protraction, suggesting adaptive motor control (**Fig. 5a**). In free air, these cycles are referred to as double-pumps^25,26^. When additional pumps occur following contact, they are called touch-induced pumps^25^ (TIPs). The underlying mechanism and functional advantages of TIPs are not completely known. However, it has been shown that TIPs are selectively gated by attention, emerging with a higher probability under attentive exploration^27^. As our results demonstrate that convergence towards α^*^guides contact-related behavior, we asked whether proximity to α^*^may form a useful lens for studying the dynamics of TIP emergence. We defined a TIP-cycle as any cycle in which a delayed protraction followed contact (**Fig. 5a**; see **Methods**). Consistent with the literature^25^, TIP-cycles exhibited amplitudes similar to those of non-TIP contact cycles, with durations increased by 38% on average (**Fig. 5b,c**; Wilcoxon rank-sum test, *p* < 10^−1^^4^). The probability of TIP emergence increased in the last cycle of the approach phase, compared to preceding cycles (**Fig. 5d**; Fisher’s exact test, *p* < 0.001). However, TIP probability was not dependent on contact cycle number relative to last cycle (logistic regression, *p* = 0.842), suggesting that TIPs do not simply become more likely as the approach phase progresses.

**Fig. 5.**
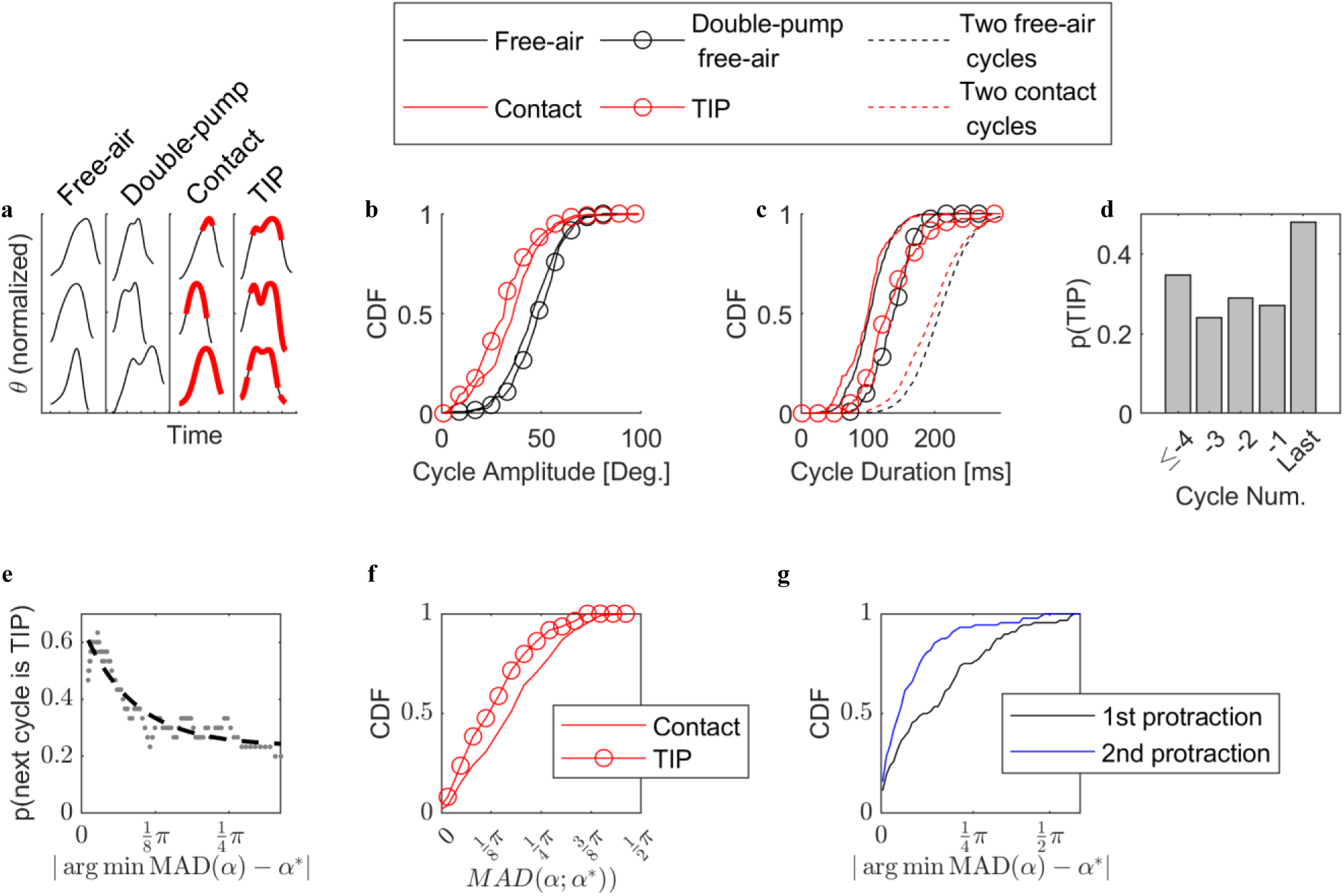
Motor-sensory plane dynamics predict touch-induced pumps and frame their functional advantage. **a.** Examples of different cycle types, from left to right: free-air, free-air double-pump, contact, touch-induced pumps (TIP-cycles). Contacts are denoted by thick red traces. TIP cycles refer to any cycle in which an additional protraction trace follows a contact. **b.** CDFs of cycle amplitudes, by type (*n*_*free*−*air*_ = 467, *n*_*contact*_ = 218, *n*_free–air double – pump_ 120, *n*_*TIP*_ = 119). In panels **b**,**c** only, data are used from both sides of the snout throughout object approach. **c.** CDFs of cycle durations, by type. TIP-cycles had longer duration than contact cycles (one-tailed Wilcoxon rank-sum test, *W* = 30,145, *p* < 10^−1^^4^). **d.** Probability of TIPs does not depend on cycle number (logistic regression, non-significant, χ^2^ = 0.039, *p* = 0.842; *n*_*cycles*_ (from last backwards): 115, 89, 52, 25, 26). However, TIPs were more likely in last cycles than non-last cycles (one-tailed Fisher’s exact test, *p* < 0.0005; *n* _*last*_ = 55/115, *n n(TIP)_non–last_* = 54/192). This and following analyses include only data from trials with a unilateral contact preference during approach. **e.** Probability of TIPs depends on the α value in the protraction of a preceding contact cycle. For each contact cycle’s protraction period, the α that minimizes MAD was obtained (total *n*_*followed by contact*_ = 74, *n*_*followed by TIP*_ = 45). The x-axis denotes absolute deviation of this value from α^*^. The y-axis denotes TIP probability in the subsequent cycle. Scatter points show data sorted along the x-axis and smoothed by a moving average with a window size of 30 observations. Dashed black trace denotes an exponential fit to the scatter points 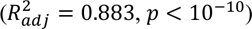. **f.** TIP-cycles clustered more tightly around α^*^ than non-TIP contact cycles, quantified via MAD (α; α*) (one-tailed Wilcoxon rank-sum test, *n*_*tip*_ = 109, *n*_*contact*_ = 198, *W* = 33,157, *p* < 0.001). **g.** For each TIP-cycle, we calculated the α that minimizes the MAD, separately for the two protraction periods. The x-axis corresponds to the absolute deviations of this value from α^*^. The second protraction’s α values were significantly closer to α^*^ than those of the first protraction (one-tailed Wilcoxon signed-rank test, *n* = 88 cycles, *W* = 2,938, *p* < 0.001).

We first examined whether our data align with schemes of attentional gating^27^, by testing whether dynamics of α predict probability of TIP emergence in following cycles. For this purpose, cycle pairs were analyzed in which the former was a non-TIP contact cycle, and the latter was either a TIP-cycle or not. This analysis showed that the more tightly α clustered around α^*^ in the protraction phase of the preceding cycle, quantified via the angle that minimizes MAD, the more likely the succeeding cycle was to be a TIP-cycle (**Fig. 5e**; exponential fit, *R*^2^ = 0.883, *p* < 10^−10^). Thus, TIPs emerged with higher probability following contacts that clustered tightly around α^*^. Given the link between α^*^proximity and the probability of TIP emergence, we next asked whether TIPs reciprocally shape α’s dynamics. We quantified tightness around α^*^ via *MAD*(α; α^*^). This analysis, which considered the full cycle, revealed that TIP-cycles clustered more tightly around α^*^than non-TIPs (**Fig. 5f**; Wilcoxon rank-sum test, *p* < 0.001). Consequently, we asked whether α dynamics evolved towards α^*^ within TIP-cycles. TIP-cycles were separated into their initial and second protraction. The second protraction was closer to α^*^ than the first, quantified via the angle that minimizes MAD **(Fig. 5g**; Wilcoxon signed-rank test, *p* < 0.001). In sum, we found that the occurrence of TIPs is related to proximity to α^*^ in preceding cycles, and that TIPs themselves are associated with expedited convergence towards α^*^ within a single cycle.

## Discussion

Detecting external objects during active exploration poses a challenge: how can perceptual confidence arise from continuously varying motor and sensory signals? We jointly tracked motor and sensory dynamics of individual contacting and non-contacting whiskers, at millisecond timescale, in freely moving rats localizing and exploring objects. We found that as rats approached an object, the non-contact distribution covered most of the motor-sensory phase-plane (*θ̇*, *κ̇*), with some preference for positive changes in base-angle (right hemi-plane), likely because protractions are typically of longer duration than retractions (**Fig. 2b**,**c**). In contrast, consecutive contacts gradually evolved to cluster around an invariant ratio between changes in base-angle and base curvature, termed α^*^. The experimental design did not aim to guide behavior towards maintaining α^*^; rats were trained to report on an object’s identity, not its presence or location. As rats moved freely in an uncertain environment, they strategically and adaptively tuned their motor-sensory interactions to allow convergence to α^*^ if an object is contacted, or divergence towards α ≈ 0 otherwise. Thus, successful convergence at α^*^ likely indicates to the rat a high probability of contact.

Is α directly involved in object detection or is it an epiphenomenon? Our results support active involvement of α. First, regardless of the number of contacts made during the approach phase, the last contact cycle was strongly associated with α being close to α^*^. Second, correlation analyses suggested that α specifically was a driving force not only for contact perception but also for head-motion. Third, proximity to α^*^ induced emergence of TIPs in following cycles. Fourth, TIPs further induced rapid convergence of α towards α^*^ within a single cycle.

Previous works in simulations, mechanical models and head-fixed rodents have shown that object location in the horizontal domain (spanned by azimuth and radial distance) can be mapped to various ratios of sensory and motor variables, most notably between specific measures of κ and *κ̇* and specific measures of θ and *θ̇*^4–6,8,28–30^. Furthermore, it has been suggested that detecting an object may require more than one contact when relying exclusively on these ratios, unless high-frequency vibrational components are specifically considered^28^. We showed here that in freely moving rats, one phase plane is preferred for detecting contacts with objects – the (*θ̇*, *κ̇*) plane. Thus, we propose that when the head is free to move, rodents prefer to base their confidence in contacting an object on comparing *κ̇* and *θ̇*. The ratio of *κ̇* to *θ̇* has previously been suggested to encode radial distance^5,7^ (RD). In our data, RD showed comparable correlations with both α and *r* (**Fig. 4**). However, *r*’s dynamics did not fully align with those of RD, and *r*’s variance increased throughout approach. Thus, α was the most plausible variable encoding RD, consistent with previous suggestions.

Our findings challenge traditional theories in which percepts are constructed via open-loop sampling of sensory information. One salient open-loop model of perception, often used to model perceptual decision making, is evidence accumulation^31^. Under open-loop evidence accumulation, percepts are often assumed to emerge from the gradual integration of sensory information over time without concurrent adaptation of exploration movements. Our main finding, that when encountering an object motor-sensory dynamics converge to a unique invariant ratio from a variety of initial conditions, is not consistent with such models. Our data show rapid reciprocal feedback between motor and sensory signals in the form of convergence towards α^*^ both across and within whisking cycles (whose typical duration is ∼50-120ms). These timescales are shorter than those typically associated with decision-making in evidence accumulation paradigms. Our results, therefore, best align with closed-loop frameworks which feature reciprocal feedback between motor and sensory components^15,32–37^. Closed-Loop Perception^2^, further predicts that the invariant ratio we outlined is more than a motor-sensory contingency^3^; it should specifically be an attractor in the motor-sensory phase-plane^36,38^.

Based on previous works, several criteria seem appropriate for characterizing attractor dynamics in behavioral data: 1) states should be localized to a low-dimensional subset of the space; 2) states should evolve back to attractors after perturbation; and 3) the set of states should demonstrate invariance, e.g., across conditions^39^. The scope of the current work does not examine external perturbations. However, patterns of convergence suggest that self-initiated perturbations, in the form of movements away from α^*^ during contact, tend to converge back to it from one direction (**Fig. 3**). In addition, we suggest that the two other criteria are also supported in our results. Namely, the invariant ratio we describe, α^*^, approximates a one-dimensional line in two-dimensional space or a specific angle in one-dimensional space, and is invariant to the object’s details (**Fig. 3f**). We thus propose our findings describe a motor-sensory invariant representing contact with external objects, and potentially – a perceptual attractor of such a contact.

## Methods

### Animals

All animal experiments were performed according to protocols approved by the Weizmann Institute’s Institutional Animal Care and Use Committee (IACUC). Data are reported from two male Wistar rats. Rats were housed in a communal animal housing facility and were introduced to the experiment room for each session. In the housing facility, rats were kept in a 12-hour light/dark cycle. Upon acquisition of the animals, at the age of 8 weeks, they were habituated to hand handling for ∼6 sessions within the experiment room. Food was available ad libitum, while water was restricted to motivate task performance. Water deprivation periods were varied based on task-performance, and eventually set at ∼16 hours per day. Deprivation was only used on days in which rats performed the task. To facilitate whisker tracking, rats’ whiskers were trimmed, except for row C, on both sides of the snout, in the four sessions that compose the data reported. Trimming was carried out under sedation, and data were never collected after sedation on the same day.

### Experimental Setup

The setup, illustrated in **Extended Fig. 4**, comprised two parts: an arena and a task area, connected by a single door. The arena’s dimensions were 30 × 22 × 30 cm (L × W × H). The task area was an open platform, which had dimensions of 23.5 × 18 cm (L × W). The entire structure was made from clear Perspex and was mounted on poles to discourage rats from straying away from the task area. Within the task area, a single object was presented at a time, using the combined operation of two E19-1 linear servo motors (ElectroCraft, Dover, NH). The arena had four doors: one was the front door, leading to the task area. The three remaining doors each restricted access to a sipper spout when they were closed (E24-01 Optical Lickometer H20-94 single photocell sensors, Coulbourn Instruments, Allentown, PA). Doors were controlled by HS-322HD Standard Heavy Duty Servo motors (HiTec, Poway, CA). Rewards in the form of juice release were controlled by a SCH284A005 two-way normally-closed pinch valve (NResearch, West Caldwell, NJ). Trial management was performed by C++ based software developed in-lab, which automatically initiated trials, chose objects to be displayed based on presentation history while controlling overall presentation probability, controlled motors for object presentation, allowed or restricted access to task area and sipper spouts, administered juice rewards, and logged trial information along with acquired video identities.

### Stimuli

Objects were three metal poles created to specification by the Weizmann Institute’s in-house workshop. Poles had a height of 7cm and a diameter of 48 mm. Poles had one of two shapes, rectangular or circular, and one of two textures, smooth or rough. Rough objects had grooves running along their circumference. Grooves were spaced horizontally by 300μ*m*, had a length of 300μ*m*, and a depth of 50μ*m*. The objects consisted of the following feature combinations: rough circular pole, smooth rectangular pole, and smooth circular pole.

### Video Acquisition

An IR backlight was positioned beneath the task area, to support video acquisition in the dark. The MB-OBL9×9-IRN-24-1V OmniLight flat dome backlight produced 830nm infrared light (Metaphase Technologies, USA). Videos were acquired using an IR-sensitive CL600×2 camera (Optronis GmbH, Germany), with a resolution of 1024 × 1280 pixels at a rate of 500 frames-per-second (FPS). Video acquisition was performed by software developed in-lab, which automatically triggered video acquisition via movement detection. A region-of-interest (ROI) was defined manually in each session, and triggering was applied when the mean difference in values of all pixels within the ROI exceeded an experimenter-determined threshold value.

### Task

After a rat was introduced to the arena, the room was darkened to eliminate visual cues. Objects and surfaces were scrubbed routinely with 70% ethanol to eliminate olfactory cues. Trials began when one of the three objects was moved into the presentation position (common to all objects in the set within a session). In three of the four analyzed sessions, the object was positioned ∼3.5 cm from the task-area door; in the fourth, it was ∼8.5 cm away. The overall duration of motor movement was kept constant for each of the three objects so their identity could not be inferred from the durations of noise produced by motor operation. The door connecting the arena and the task area would then open, allowing subjects to approach the object. As a rat entered the area, it triggered video acquisition by moving through an ROI, and as it moved back out through the ROI, video acquisition was stopped. Upon re-entering the arena, all sipper doors opened, allowing the rat to report object identity by choosing the associated sipper. A correct report was rewarded with the dispensing of ∼0.1 ml of juice. For incorrect responses, no reward was dispensed. This marked trial termination, at which point all doors were closed and trial data were logged.

### Data inclusion

Rats performed four sessions with the whisker-pad trimmed to row C exclusively. Videos from these sessions were examined by three independent annotators, and a dataset of *n* = 155 videos was used as the basis of all analyses reported here. The dataset included videos which satisfied the following requirements: acquisition was executed correctly; the video was the first (or only) one acquired in a given trial, meaning repeat excursions within a single trial were excluded; object contact had been made; and approach phase was well defined. Trials were removed only if all annotators agreed that not all criteria had been satisfied. The work discussed here is focused on the modulation of motor-sensory behavior supporting localization during object-approach. Thus, analyses presented throughout are not contingent on sipper-approach responses.

### Data preprocessing

#### Whisker tracing

Whisker tracing was performed via Whisk^40^. Snout contour tracing was performed via BioTact^41^. Inputs of head-center and nose-tip coordinates, required as input to BioTact, were extracted via an instance of the DeepLabCut network trained specifically for this purpose^42^. The head-center coordinate tracked by Biotact was used in the calculation of approach velocity (see **Data Processing** subsection). In each frame, whiskers whose distance from the snout contour exceeded 7 pixels (0.82mm) were dismissed. Snout length along the rostral-caudal axis was z-scored per video, and all whiskers whose follicle was rostral to the snout position with z-score exceeding 2 were removed as potential false-positives. Occasionally, whisker traces extended beyond the snout contour, in which case traces were trimmed to the point of intersection; for this purpose, snout traces were linearly up-sampled. For each whisker trace, a cubic spline was fitted, with piecewise third-degree polynomials of the form *Ax*^3^ + *Bx*^2^ + *Cx* + *D*. Spline knots were equally spaced at distances of 40 pixels (4.66mm) along the whisker trace. A spline fit was considered valid if the distance between all points in the original trace and the spline did not exceed 3 pixels (0.34mm), and the overall root-mean-square error (RMSE) did not exceed 0.5 pixels. Whisker traces for which the fit was invalid were rotated iteratively, in an attempt to resolve numerical issues. If no valid fit was achieved over rotations, the whisker trace was omitted from the data. The polynomial fit at the whisker’s base (close to the snout) was used to calculate base-angle θ and base-curvature κ as in^4^, using the following formulae:

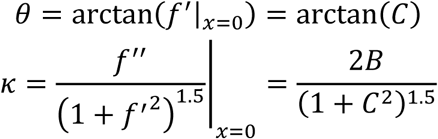

κ is inversely related to the radius of an osculating circle – i.e. a circle that best approximates the local curve at the point of measurement. A perfectly flat whisker would have an infinitely large osculating circle, and thus its corresponding value of κ would be zero. As bending increases at a point, the size of the osculating circle decreases, and κ consequently increases. Consistent with previous work^4^, curvature is expressed in units of *m*^−1^.

#### Whisker tracking

Association of whisker traces into tracks across consecutive frames was performed using code developed in-lab. The approach was Kalman-Filter-based: each whisker was represented by two pairs of (*x, y*) coordinates – at distances of 10 and 20 pixels away from the follicle (1.17 and 2.33 mm, respectively). Thus, a state-vector was assigned for each whisker trace, as an eight-dimensional vector comprising these coordinates and their associated velocities (initially set at zero). Using a simple position-velocity Kalman-Filter, predictions were made for each state-vector for a single time-step (one video-frame, 2ms). Matrices H,R,P were initialized as proportional to the identity matrix, and assumed an independent relationship between velocities (i.e. for any position-value *x*, *x*(*t* + 1) = *x* + *ẋ*). The model’s predictions were compared with measurements of whiskers in the following frame. Probabilities of assignment between each measurement and each prediction were given by the Gaussian uncertainty around the predictions. Assignment costs between prediction and measurement were then defined as the inverse of that probability. Costs that exceeded a maximal threshold were marked unassignable and omitted as potential matches. Costs were then used to resolve the assignment problem via the Munkres algorithm. Propagating the state-vector across frames allowed for formation of initial tracks from concatenation of matched pairs. After an initial track-formation step, the procedure was repeated, this time feeding each formed track into the Kalman-Filter, and searching for a prediction that has not been previously assigned anywhere within a larger temporal range of up to 4 frames (8ms). Upon completion of the second step, the procedure was repeated by propagating state-vectors backwards in time.

Note that tracks are not labeled, i.e. they are not assigned with unique identities that can identify a whisker across videos - e.g. whisker C3. This is because of slight differences in whisker-trimming, and the fact that whiskers were not always traced to full length. Instead, track membership merely denotes that all traces likely correspond to a single whisker; within the same video, several tracks may belong to the same whisker. Typically, a large portion of whisker traces were assigned to a small number of tracks, which spanned tens to hundreds of frames. Visual examination of resultant base-angle dynamics demonstrated adequate separation of whiskers, see **Extended Fig. 5**.

#### Filtering, outlier exclusion and derivative estimation

Pixel to centimeter correspondences were estimated from videos based on known landmarks. Whisker tracks were excluded from the dataset if the whisker’s median length was smaller than 5mm, or if the track’s length was shorter than 30 frames. Additionally, as κ distributions had pronounced tails, unrealistically extreme outliers were removed to reduce their effect on subsequent lowpass filtering (|κ| > 300*m*^−1^). Both θ and κ were lowpass filtered using a 1^st^ order Butterworth filter with a cutoff frequency at 200Hz. First-order forward difference method for numerical differentiation was used to estimate time-derivatives *θ̇* and *κ̇*.

## Data processing

### Threshold estimations and data transformations

For the analyses of contact and non-contact visit-rates on the (*θ̇*, *κ̇*) plane, data from frames in which the snout was in contact with the object were excluded, using a distance threshold of 1mm. The whisker-object contact threshold was determined by analyzing the log-scale distribution of whisker-object distances and identifying the inflection point between two modes, yielding a threshold of 1.34 mm.

To compute α, the (*θ̇*, *κ̇*) plane was normalized by scaling κ, to account for difference in units between *θ̇* and *κ̇*. A data-driven coefficient *c* was estimated to make the non-contact α distribution as uniform as possible, resulting in *c* = 0.48. This was performed by iterating over values of *c*, and evaluating the sum of deviations between the empirical CDF and a uniform CDF over the same range to obtain the minima. The parameter α^*^ was estimated as the α value corresponding to the peak near the positive mode of the α distribution for whisker-object contact. The resultant value was α^*^ = 1.435 ± π, and is used throughout (i.e. α^*^ is always considered equivalent to itself modulo π).

Several analyses are confined to the approach phase of each trial, defined as the period from video acquisition onset until the first snout-object contact (if one occurred). If no snout-contact occurred, the approach phase ended when head velocity along the y-axis fell below a data-driven threshold. This threshold was estimated by filtering the head-center y-position with a 1st-order lowpass Butterworth filter (100 Hz cutoff), then fitting a three-component Gaussian mixture model to the velocity distribution across all data. The mean of the leftmost component was set as the threshold, resulting in 0.79 cm/s. Using this threshold, approach velocity traces aligned well with one another, demonstrating a stereotypical trajectory (**Extended Fig. 6**).

For most cycle-based analyses, data occurring after the approach-phase were excluded. However, for analyses centered on α dynamics during touch-induced pump cycles (TIP-cycles), the entirety of data from the last-contact cycle of the approach phase was used, even if it extended beyond the approach phase. This is because the last cycle of the approach phase had a higher probability of TIP emergence compared to preceding cycles, and thus including the full cycle ensured that the second protraction of the TIP was not omitted. The median number of frames added by this extension was 27, corresponding to 54ms.

### Defining whisking cycles

Whisking cycles were defined separately for each side of the snout. Per side, the median base-angle was estimated by differentiating θ across all tracks, taking their median, and then recalculating the cumulative sum. This ensured robustness to frame-to-frame variations in whisker counts. Missing data were linearly interpolated. The median angle was lowpass-filtered using a 6^th^ order Butterworth filter with a cutoff at 80Hz^25^. The lowpass-filtered signal was then high-pass-filtered using a 6^th^ order Butterworth filter with a cutoff at 7Hz (resulting in a 12^th^ order composite bandpass filter). A Hilbert transform was applied to the resultant signal to obtain an instantaneous phase^43^. Cycle boundaries were identified as peaks in the Hilbert phase near π, and adjusted to align with peaks and troughs of the low-pass-filtered signal where needed. Up and down traces were classified by the sign of the lowpass-filtered median angle’s derivative. Cycles with multiple up-traces were classified as multiple-pump cycles. A multiple-pump cycle was labeled a TIP-cycle if any whisker made contact with the object prior to the second up-trace.

### Cycle-based parameter estimation

Head movement was quantified by approach velocity, defined as the per-cycle mean of frame-to-frame differences in head–object distance along the y-axis, multiplied by the sampling rate to convert to units per second (i.e. *mean*(Δ_*t*_|*head* − *enter*_*y*_ − *object*_*y*_| × 500)), after lowpass-filtering the head’s trajectory with a cutoff at 100Hz. For whiskers, α was parameterized using two approaches, each capturing distinct aspects of its dynamics. Both estimates were computed throughout and were typically in agreement; in each analysis, the reported estimate was selected based on clarity and interpretability. The first estimate, used to establish the characteristic value of α^*^, was defined as:

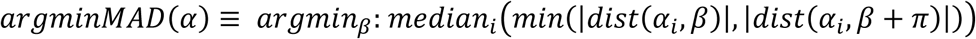

where the *dist* function denotes angular distances on the unit-circle, and α_*i*_ denotes the relevant subset of observations.

Second, a similar approach was used to calculate dispersion with respect to α^*^, calculated explicitly as:

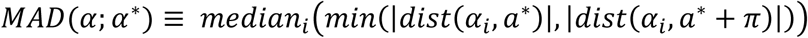

with α^*^ = 1.435 ± π, as noted above. For plane radius *r*, denoting the magnitude of the (*θ̇*, *κ̇*) signal after scaling *κ̇* and dividing by the 500 Hz sampling interval, cycles were parameterized using the 90^th^ percentile, as trajectories typically overlapped, and differed mostly by their peaks. Finally, for contact observations, radial distance from whisker base to contact point, denoted RD, was parameterized per cycle as the median of all contact observations. For the variables α and *r*, cycle*-*based analyses typically included cycles from the approach phase of trials in which 90% of whisker-object contact observations were unilateral, starting at first contact. The label ‘contact’ denotes whiskers on the contacting side during actual object contact, the label ‘same side’ denotes non-contact observations on the contacting side, and the label ‘other side’ denotes non-contact observations from overlapping cycles on the opposite side. For the distributions of TIP-cycle duration and amplitude, data were aggregated from both sides of the snout, but still only from the approach phase.

In two TIP-related analyses contingent on α, only observations occurring during protraction periods were used (i.e. up-traces of the whisking cycle). In these cases, estimates of *argminMAD*(α) still considered both the 1^st^ and 3^rd^ quadrant sections of α^*^, as flow analyses have demonstrated that contact dynamics during protractions may flow to either of them.

### Kernel-smoothed function reconstructions

For the analyses focused on flow of α dynamics, smooth reconstructions of functions were derived from data using kernel-smoothing. This approach avoids reliance on bin size, instead weighting all observations by their distance from the point of estimation. Reconstructions were performed separately for contact and non-contact observations during approach, estimating two functions: mobility and directional tendency. Both functions relate Δα (the change in α over 4 ms) to the current value α. Large steps were excluded 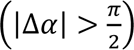. Mobility at α = *x* was estimated as the kernel-weighted median of |Δα| values near *x*, using Gaussian kernel weighting with bandwidth σ = 0.15. Directional tendency at α = *x* was defined as the difference between the kernel-weighted fractions of positive and negative Δα values, using the same kernel and approach.

### Data usage and sample sizes across analyses

The initial contact probability analyses used all trials, *n*_*trials*_ = 155, excluding frames without snout-object contact (with *n*_*frames*_remaining 83,701/94,210). For all other analyses, only data from the approach-phase is used, *n*_*whiskers*×*frames*_ = 219,346. Cycle-based analyses included trials where ≥90% of contact observations during approach were from one side, *n*_*trials*_ = 83. ^These included: *n*^*first cycles* ^= *n*^*last cycles* ^= 83; *n*^*interim* (*same side*) ^= 97;^ *n*_*interim (other side)*_ = 102. When trials with a single contact cycle were included, this added *n*_*trials*_ = *n*_*cycles*_ = 26. Thus, total *n*_contact *cycles*_ = 289. Cycle correlation analyses therefore comprise *n*_*cycles*_ = 289, and those using cycle lagging comprise *n*_*cycles*_ = 289 − 83 − 26 = 180.

Characteristic amplitude and duration of single/double-pump cycles were obtained from all whisking cycles bilaterally from trial onset throughout the approach phase (*n*_*trials*_ = 155), yielding: *n*_*free*−*air*_ = 467, *n*_*contact*_ = 218, *n*_*free*−*air double*−*pump*_ = 120, *n*_*TIP*_ = 119. All other TIP-related analyses used unilateral contact approaches (as in previous analyses); however, these analyses included data until the end of the last whisking cycle, even if it extended past approach termination. Subsequently, *n*_*trials*_ = 115, *n*_*TIP*−*cycles*_ = 109, *n*_*contact*−*cycles*_ = 198. For the analyses of TIP emergence probability, *n*_contact *followed by TIP*_ = 45, *n_contact followed by contact_* = 74. For analyzing within-TIP dynamics across first and second protractions, *n*_*TIP*−*cycles*_ = 88; cycles were excluded where contact was absent in either protraction, or occurred in tracks that were removed from analyses (see *Filtering, outlier exclusion and derivative estimation*).

## Statistics

Statistical tests are two-tailed unless otherwise specified. To minimize assumptions about properties of data distributions, Wilcoxon rank-sum test was used for comparing central tendencies of two independent samples, and Wilcoxon signed-rank test was used for comparing paired samples or a single sample and a known value. A Brown-Forsythe test was used to test for equality of variances. Multiple comparisons were corrected using false-discovery rate thresholding (FDR) where applicable. Significance was thresholded at *p* < 0.05. Pearson’s correlation coefficient was used for correlation analyses, and differences between correlations were tested via Fisher r-to-z transformation. Fisher’s exact test was used to compare binomial outcomes between two samples. When discussing entropy, Shannon entropy was computed using 360 bins over the range [−π, π]. All physical distances (e.g., whisker-object, snout-object, etc.) refer to two-dimensional Euclidean distances. All distances relating to α, e.g. deviations in *MAD*(α; α^*^), refer to absolute angle difference on the unit-circle. For analyses of TIP-cycle and free-air double-pump duration, the distribution of two complete cycles was obtained by generating all possible pairs of single-pump cycles from the corresponding populations.

Permutation tests were used to estimate significance in five cases: 1) for entropy differences of the α distribution in contact and non-contact; 2) for differences in Jensen-Shannon divergence between two pairs of two-dimensional (α, *r*) distributions; 3) for comparing *argminMAD*(α) obtained from different object groups by shape or texture, preceded by an omnibus test that compared the deviation of each object’s median α from the grand median across all labels; 4) for comparing differences between sample medians when group size was small (*n* = 26), in the analysis comparing single-contact cycles with first or last cycles; 5) for assessing the relationship between head position and α across cycle pairs (**Extended Fig. 2**), using shuffled pairs to generate the null distribution. In each case, observations were randomly shuffled to create control groups with the same size as the original groups, and the relevant statistic (difference in entropy, JSD, α, or median) was recalculated. This process was repeated 1,000 times to generate a null distribution, against which the observed statistic was evaluated for significance.

Similarly, bootstrapping was used to estimate confidence intervals by resampling each of the contact and non-contact groups with replacement 100 times and repeating the kernel-smoothing function estimation procedure. Confidence interval bootstrapping was employed for the mobility and directional tendency functions of the flow analyses. Resampling was performed twice: once exclusively within the contact group (to generate the error bars associated with contact), and once across the combined contact and non-contact groups, resampling to match the size of the contact group (labeled ‘bootstrapped’ and shown in gray). Bootstrapping was similarly used to obtain confidence intervals for the flow function in the extended figure associated with this analysis.

Where exponential models were fit to the data, parameters were estimated using nonlinear least squares optimization. Model performance was quantified using *R*^2^, and statistical significance was computed using an F-test comparing the full model to a null, intercept-only, model. Similarly, where logistic regression was used, statistical significance was computed using a χ^2^-test comparing the full model to a null model.

## External codes

In this work, we relied on several existing codes from MATLAB Central File Exchange. These include: ‘Circular Statistics Toolbox’ by Philipp Berens; ‘Generate maximally perceptually-distinct colors’ by Tim Holy; ‘fdr_bh’ by David Groppe; ‘inpaint_nans’ by John D’Errico. Additionally, the following are used in the pre-processing pipeline generating the analyzed data: ‘Munkres Assignment Algorithm’ by Yi Cao; ‘SLM - Shape Language Modeling’ by John D’Errico; ‘Fast and Robust Curve Intersections’ by Douglas Schwarz; ‘curvspace’ by Yo Fukushima. These codes are linked within the codebase.

## Code and data availability

Data and custom code are available on request from the corresponding authors.

## Contributions

G.N. conceived and designed the study, conducted the experiments, developed whisker tracking and analysis programs, analyzed the data, interpreted the results and wrote the paper. I.S-S. designed and supervised data analysis, interpreted the results and wrote the paper. E.A. Conceived and designed the study, supervised the experiments and data analysis, interpreted the results and wrote the paper.

## Acknowledgements

We thank Mitra Hartmann, Mathew Diamond, Daniel Shulz, Asif Ghazanfar, Ábel Ságodi, Alon Rubin and Luka Gantar for their invaluable feedback and insightful comments on earlier drafts of the manuscript.

This project has received funding from the European Research Council (ERC) under the EU Horizon 2020 Research and Innovation Programme (grant No. 786949), the United States-Israel Binational Science Foundation (BSF, grant No. 2021327), the Israel Science Foundation (ISF, grant No. 2237/20), the Weizmann-UK collaboration grant and a research grant from the Estate of Thomas Gruen.

---

**Extended figure 1.**
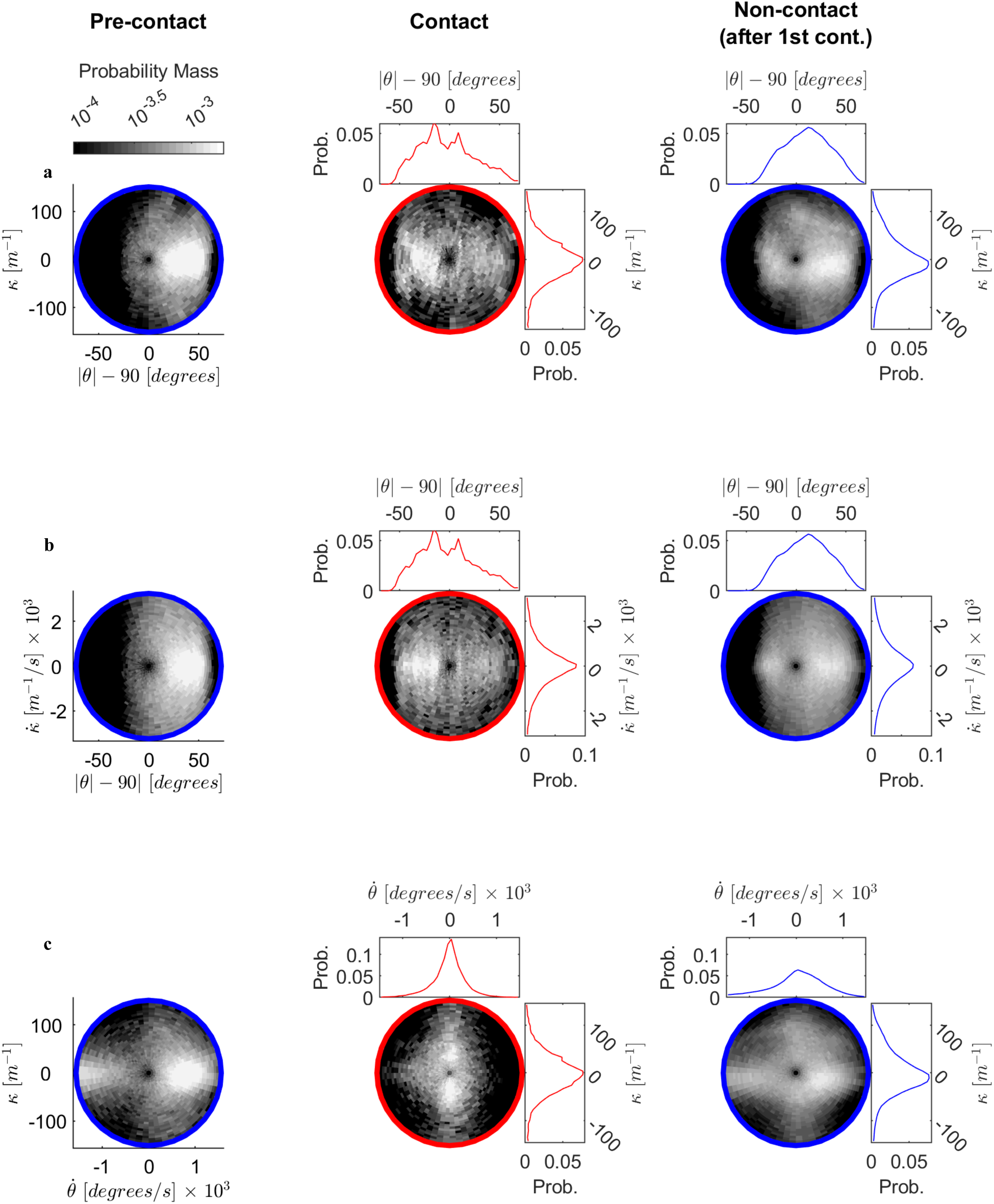
Alternative motor-sensory phase-planes for contact. Each panel depicts the phase-plane associated with a different combination of motor-sensory variables, in correspondence with Fig. 2. **a.** (*θ*, κ) plane. Sample sizes are slightly larger than those of derivatives-based planes, which require frame-to-frame matching (*n*_*pre* −contact_ = 124,479, *n*_*contact*_ = 46,011, *n*_*non − contact*_ = 464,410). **b.** (*θ*, *κ̇*) plane. **c**. ( (*θ̇*, *κ̇*) plane. For panels b and c, sample sizes are as in Fig. 2.

**Extended figure 2.**
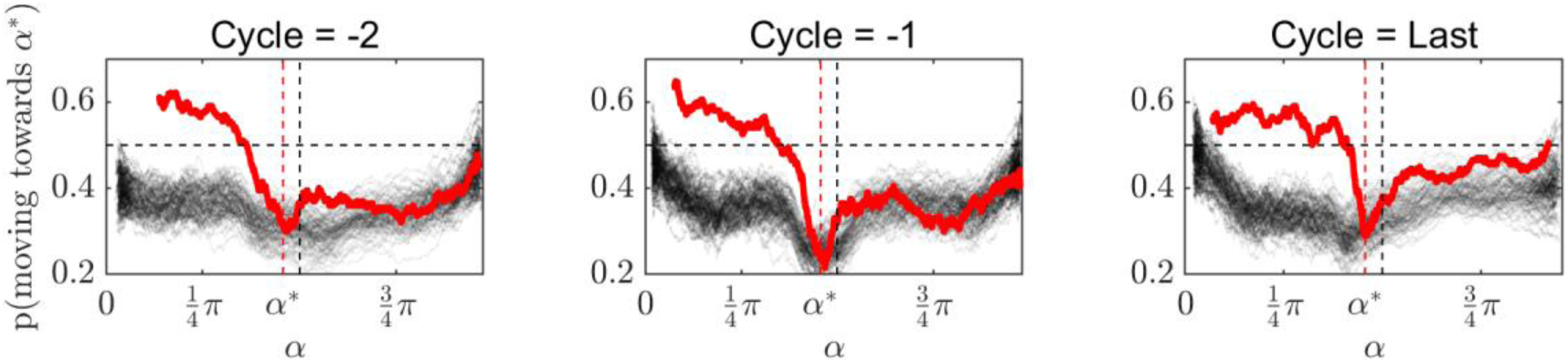
Unidirectional flow toward α* within contact cycles. The flow function employed here assigns a binary Fig. 3a**,b**. A cycle-based approach was used, as in Fig. 3c**,d**. Within each cycle, both the sorted α values and their corresponding binary labels were smoothed using a moving average with a window size of 200 samples. Red traces represent contact observations. Black traces represent bootstrapped null distributions generated from 100 resamples of the combined contact and non-contact data (with replacement), each resample matching the number of contact observations per cycle. Sample size for each trace: *n*_*whiskers* × *frames*_ = 2315, 3156, 1838, for cycles = *last*, −1,−2, respectively. The flow function was estimated in the range [0, π]. This analysis supports a unidirectional flow toward α^*^ during contact, but not during non-contact, consistent with results in Fig. 3. Moreover, it reveals that this flow pattern is stable across cycles.

**Extended figure 3.**
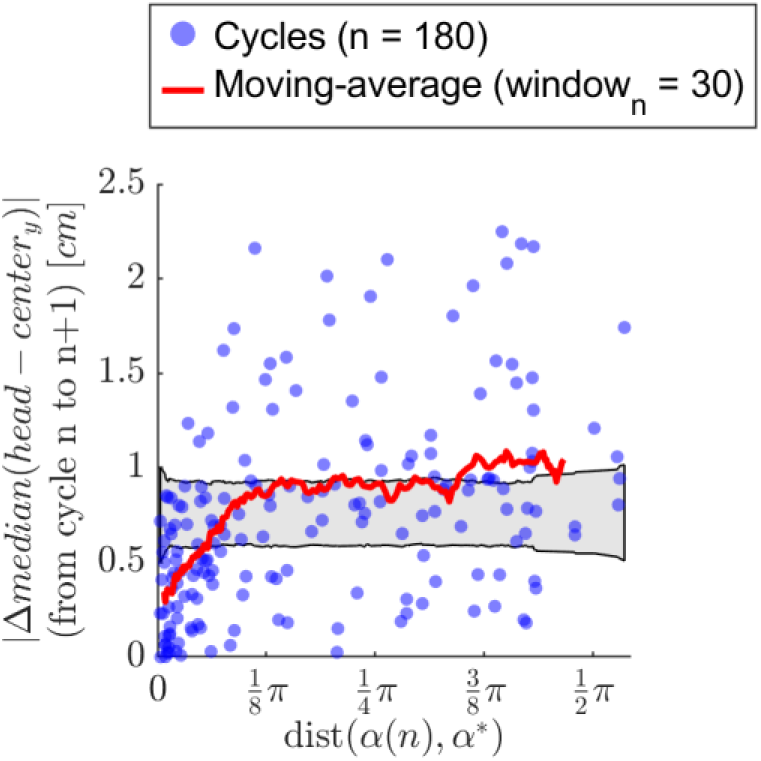
Head-center position changes less across cycles as α approaches α^*^. The absolute change in median head-center position along the y-axis was calculated between consecutive cycles. This change is plotted against α’s distance from α^*^ in the first cycle of the pair. This analysis affirms the relationship observed in Fig. 4 between α^*^and approach velocity: as α nears α^*^, changes decrease in the following cycle. Note that last cycles always occur as the second cycle in each consecutive pair. Therefore, the relationship between α and head movement captures predictive dynamics rather than simply reflecting their simultaneous convergence at the end of the approach phase.

**Extended figure 4.**
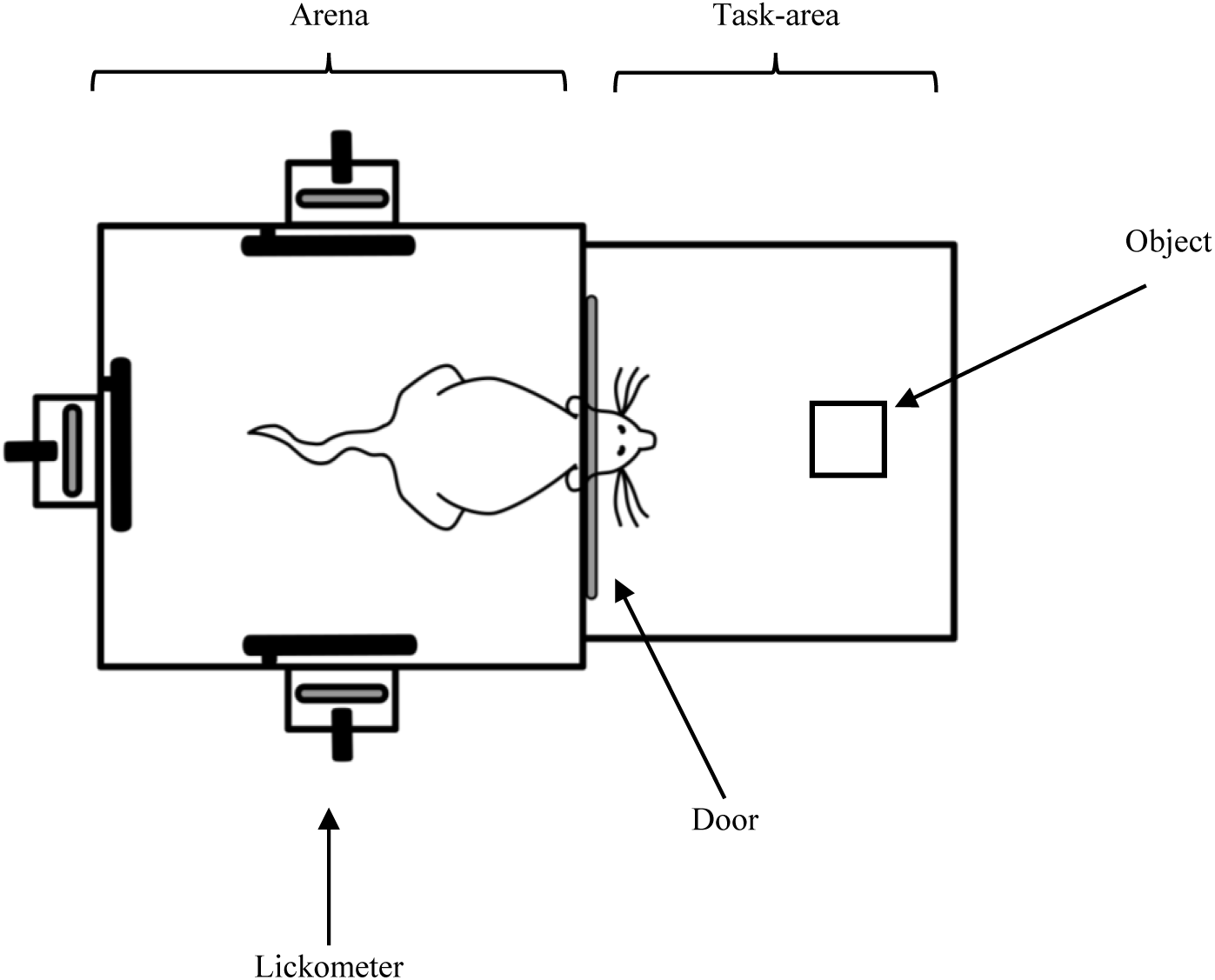
Experimental setup. Schematic illustration of the arena used in the experiment.

**Extended figure 5.**
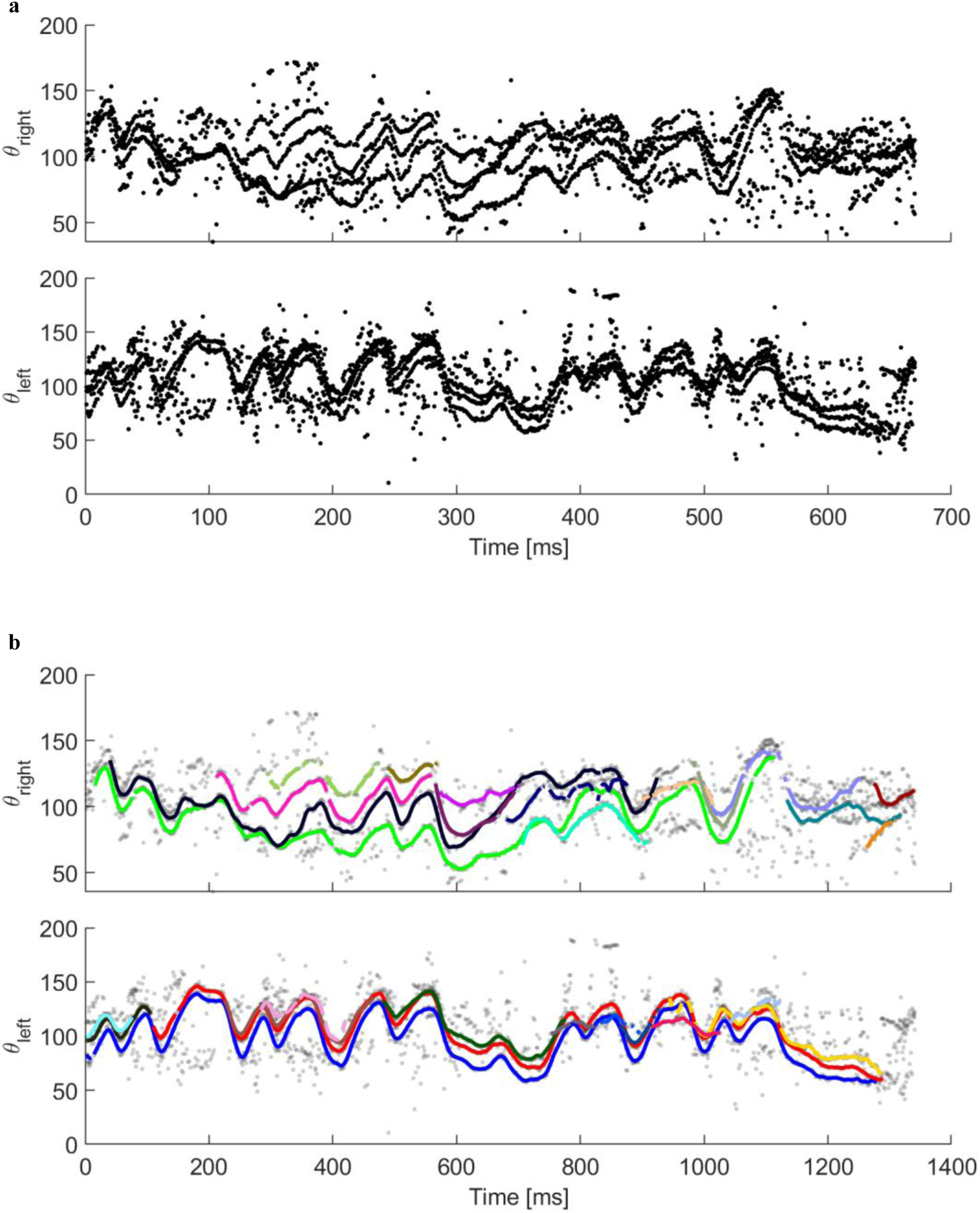
Example of whisker tracking. **a.** Angle measurements for all whisker traces, before tracking, for each side of the snout, from a single trial. **b.** Gray markers denote the same data as in **a**. Colored traces represent smoothed angles from the 35 longest tracks, with each color corresponding to a unique track.

**Extended figure 6.**
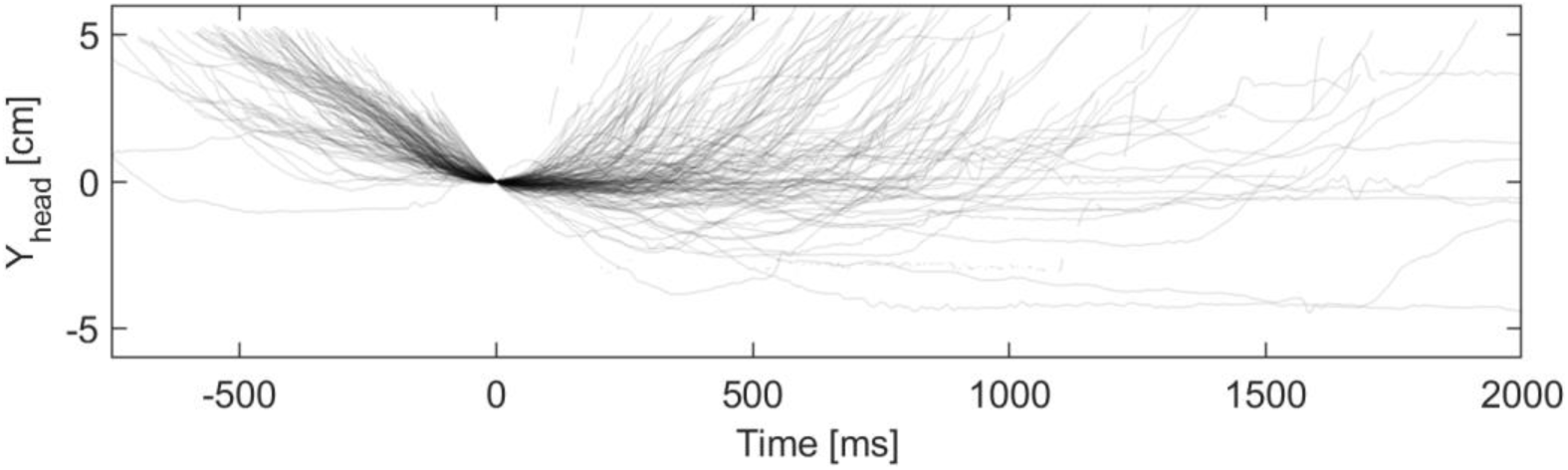
Head trajectories follow a stereotypical pattern during object approach. Traces of head position along the video’s y-axis across trials. Time and head position are aligned to zero at the end of the approach phase. Traces before *t* = 0 reveal a consistent pattern during approach.

